# Pseudo-data generation allows the statistical re-evaluation of toxicological bioassays based on summary statistics

**DOI:** 10.1101/810408

**Authors:** Ludwig A. Hothorn, Felix M. Kluxen, Mario Hasler

## Abstract

Sometimes a re-analysis of toxicological data is useful. However, this usually requires the availability of the original data and in many cases only summary data are available in the publications. Here the generation of pseudo-data under certain assumptions using extension packages in the open-source project R on statistical computing is shown. Several case studies are used to illustrate the applicability in regulatory toxicology.

## 1 Introduction

Results in publications on toxicological bioassays are usually presented as summary statistics; often in bar chart plots or table format, showing means, standard deviations and sample sizes. This practice has been repeatedly criticized (6; 2; 21) because potential sub-grouping in the data is hidden and the validity of the performed statistical analysis cannot be peer-reviewed. In internationally accepted test guidelines for regulatory bioassays, the reporting of individual data is mandatory, but often not publicly available, because they may support commercial interests. However, summary data usually are available, e.g. from the European Chemicals Agency (ECHA) or the European Food Safety Authority (EFSA), depending on the regulatory field.

When data are presented as means and standard deviations, the authors stating such values implicitly assume normal distribution of the reported data, irrespective of the actual underlying empirical distribution of the data. Therefore, it is possible to generate individual observations, here denoted as pseudo-data, from these summary statistics using a simulation algorithm.

Pseudo-data can be used for peer-review, re-analysis by different or novel methods, educational purposes and for incorporating multiple data sets in an analysis, when only summary statistics are available. While the original data are obviously more valuable, the pseudo-data approach can be a compromise; e.g. when individual data is hard to obtain or not electronically available - the latter is sometimes the case for historical control data.

We demonstrate the use of the simulation algorithm in the function `ermvnorm` within the *SimComp* package, available for the open-source statistical software R. This function is based on the function `rmvnorm` of the package *mvtnorm*. This method differs from other random data generation algorithms in that a multivariate normal distribution with an exact nominal mean vector, exact variance vector and approximate correlation matrix (for multiple endpoints or repeated measures) is calculated. (Note that random numbers with exact mean and standard deviation cannot be used for simulations concerning type I error or power of statistical tests.)

Multiple case studies are presented and an appropriate re-analysis is demonstrated: i) repeatability by a second assay, ii) fold change effect size, iii) taking variance heterogeneity into account, iv) considering dose as quantitative covariate and/or qualitative factor, v) user-defined contrasts, and vi) multiple endpoints. In a seventh case study the limitation of the approach is shown because the normal distribution assumption failed.

The presented method can be easily applied by statisticians and toxicologists by using the example R-code provided within this manuscript.

## 2 Generating random numbers with exact mean and standard deviation

First, summary statistics are selected, e.g. from a publication table for 7 antioxidant indices in epididymides of mice exposed to 0 (control), 25 (dose1), 50 (dose2) and 100 (dose3) mg/L sodium fluoride as shown by the following table from (19). Second, sample sizes (n), arithmetic means (mean) and standard errors (sd) for a particular endpoint, for example the low total antioxidant capacity (T-AOC), are simply transferred to the following R code:

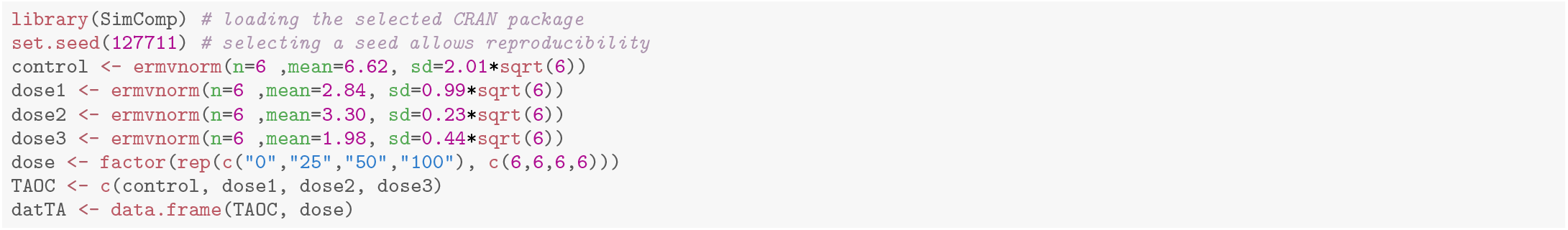

**Figure 1:**
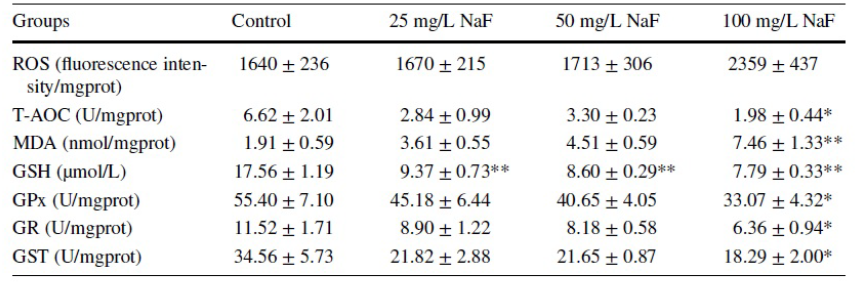
Table: Summary data table in Sun et al., 2018

The object `TAOC` contains the pseudo-data with the qualitative factor `dose` and can be used for re-analysis. These data can be easily visualized by modified box plots using the R-package `toxbox` (13)

**Figure 2:**
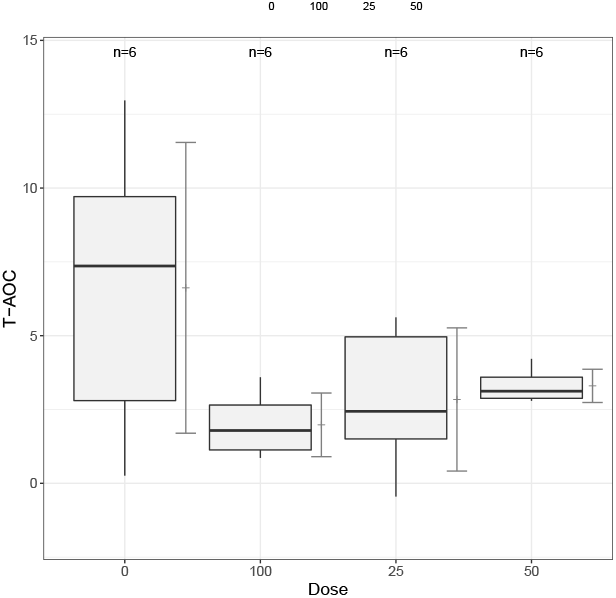
Box plots of T-AOC pseudo-data.

An extension of this example considering multiple correlated endpoints and repeated measures for assumed or random correlations is available and is demonstrated in the sixth case study below.

The simple idea behind the function `ermvnorm` is to apply the function `rmvnorm` of the package *mvtnorm*. This function generates *n* − 2 random vectors from a *k*-dimensional multivariate normal distribution whose mean vector is given by `mean` and corresponding vector of standard deviations (SD) by `sd`. If only one endpoint is of interest, dimension *k* equals 1. For more details, see (14). These *n* − 2 random vectors do not yet have the indented exact mean and SD. However, based on the first *n* − 2 vectors, a system of equations is formed by the usual formulas of sample mean and sample variance. The remaining 2 vectors are obtained as solution so that the specified values for the mean vector `mean` and for the vector of standard deviations sd are achieved. Depending on the first *n* − 2 random vectors, the underlying system of equations may have no solution. In this case, `ermvnorm` creates a new set of *n* − 2 random vectors until a valid pseudo-data set is obtained or until the maximum number of tries is reached.

## 3 Case studies

The following case studies show the applicability of the method for re-analysis for several common statistical scenarios encountered in toxicology.

### 3.1 Demonstrating repeatability

Demonstrating repeatability of a previously observed effect appears relatively simple, e.g. by a second assay under comparable conditions and seems necessary in the current reproducibility crisis. (1) argue that this might be due insufficient statistical expertise, i.e. sampling effects. Usually, data of a new assay will be compared with data from a previous assay in the literature. In this example, data for the first and the second assay were provided within the same project for the hematology endpoint mean corpuscular volume (MCV) for mice that were treated with five concentrations of sodium dichromate dihydrate (18). Summary statistics were extracted and pseudo-data generated. The boxplots in Figure 3 reveal a rather similar pattern for both studies (parallel vertical dotted lines: left study 1, right study 2) with a general tendency to lower values in the second study.

**Figure 3:**
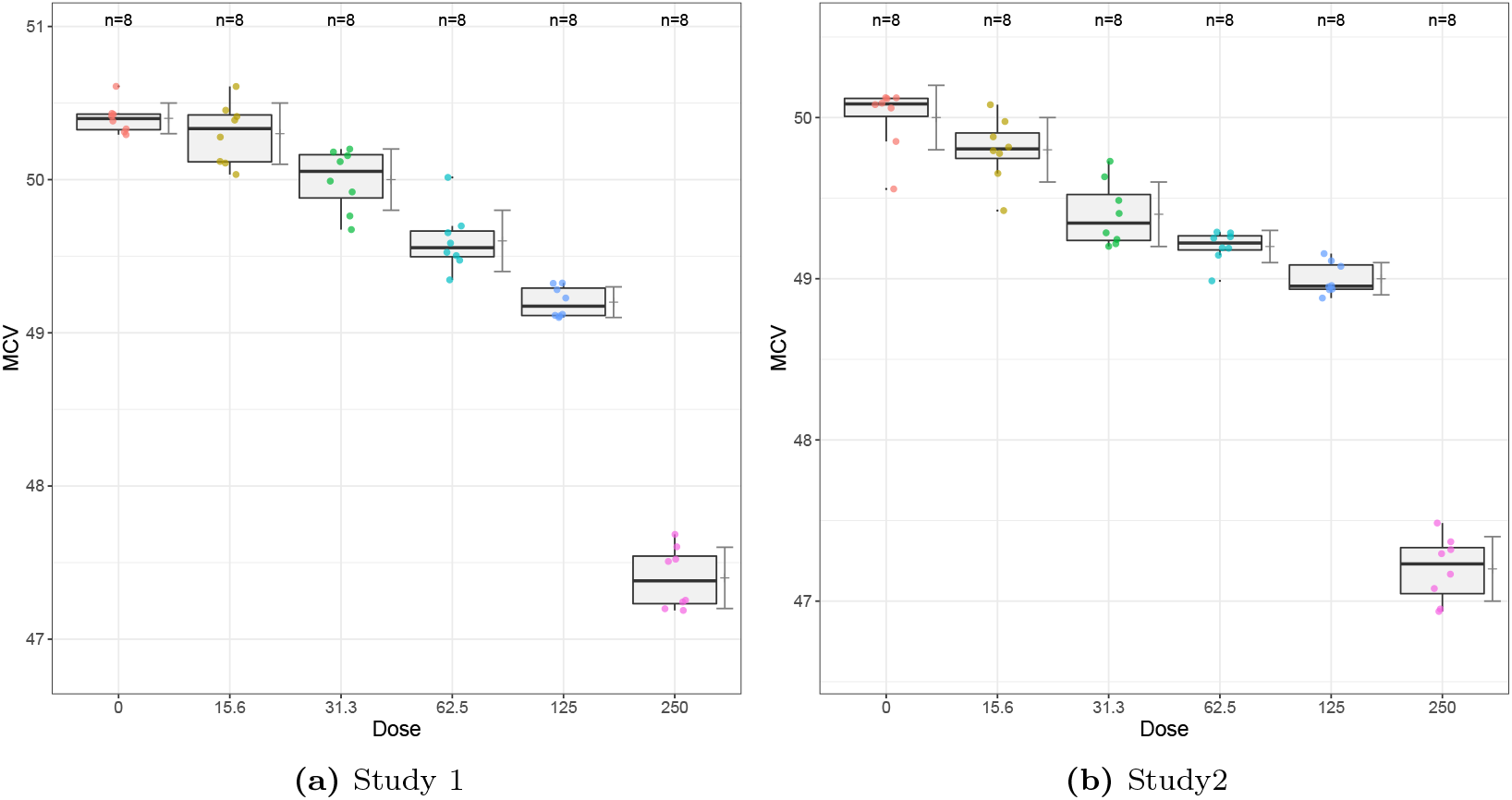
Box-plots MCV pseudo data

Table 1 shows similar effect sizes (Effect size 1, Effect size 2), i.e. the mean difference *µ*_*i*_ − *µ*_0_, and similar tiny *p*-values (*p*-value 1, *p*-value 2) for almost all comparisons against control for both study 1 and study 2, which is a convincing empirical criterion for repeatability. Irrespective of the observed additive shift, the patterns of effect sizes and *p*-values are rather similar in both studies. Alternatively, one can use a more complex model, a mixed effects model with a random factor between both studies. Again, the effect size (Effect size mixed) and the *p*-value (*p*-value mixed) are quite similar.

**Table 1:**
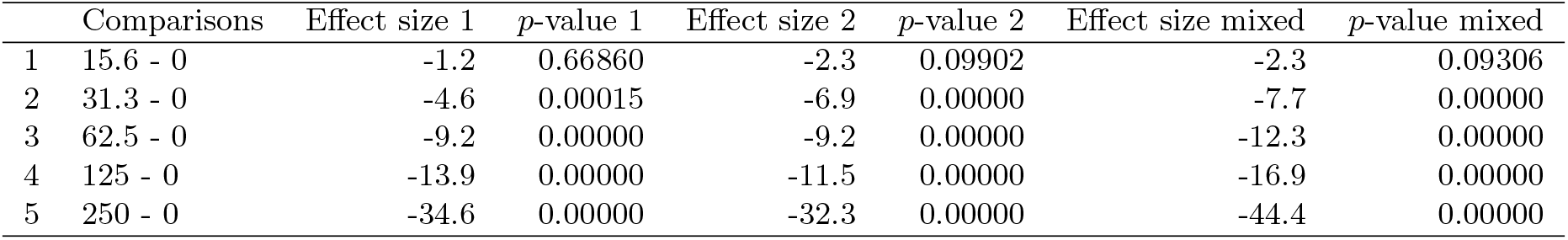
Effect sizes and *p*-values for Dunnett tests for MCV pseudo data for assay 1 and assay 2 and their joint effect size

A third and quite different approach is the claim of no interactions as a criterion for repeatability using an interaction contrast test (10). The dose-by-study interaction contrasts are estimated in Table 2 revealing quite different slopes for 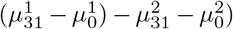 and 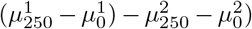, i.e. a substantial higher sensitivity of study 2 with respect to study 1. Hereby it is difficult to conclude on similarity of the two studies, as this is injured evenly with two dose comparisons substantially.

**Table 2:** Effect sizes and *p*-values for interaction contrasts for MCV data

| | Interaction contrast | Effect size | $p$-value |
| --- | --- | --- | --- |
| 1 | $((15.6 - 0):2) - ((15.6 - 0):1)$ | -0.30 | 0.017 |
| 2 | $((31.3 - 0):2) - ((31.3 - 0):1)$ | -1.70 | 0.000 |
| 3 | $((62.5 - 0):2) - ((62.5 - 0):1)$ | -0.10 | 0.781 |
| 4 | $((125 - 0):2) - ((125 - 0):1)$ | -0.10 | 0.781 |
| 5 | $((250 - 0):2) - ((250 - 0):1)$ | -1.70 | 0.000 |

### 3.2 Fold change as effect size

The majority of statistical significance tests use (implicitly) the difference of means as effect size. However, the ratio of means, denoted as fold change, is used in toxicology for specific interpretations, such as the 2-fold rule in the Ames assay (4). Unfortunately, this multiplicative effect size is sometimes used together with *t*- or Wilcoxon test for an additive effect size, e.g. for urine excretion in rats treated with seven doses of hexachloro-butadiene (20)(in their Table 5, see the simulated data in Table 3). The common approach to switch into the multiplicative model is the log-transformation (which requires the validity of a log-normal distribution, see e.g. (16)), but for normal distributed endpoints a modified Dunnett or Williams test for ratios-to-control is available (9) as an alternative.

**Table 3:**
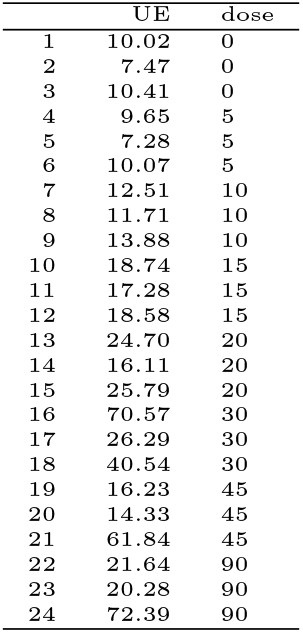
Simulated urine excretion data

The actual data from (20) is reproduced in Table 3 and the generated pseudo-data depicted in Figure 4.

**Figure 4:**
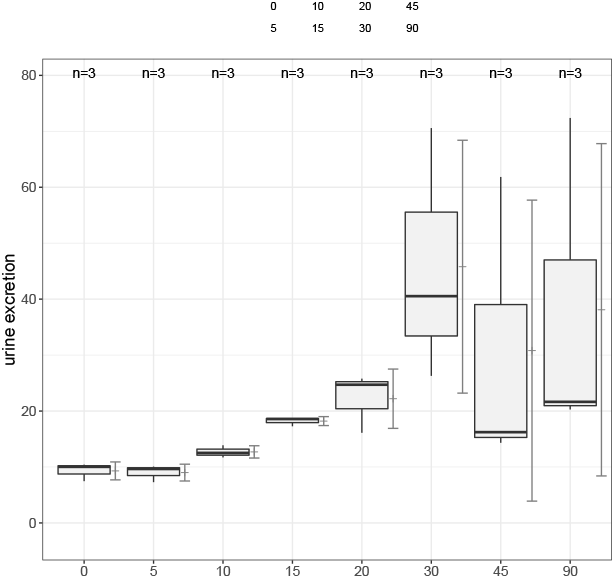
Box plot urine excretion pseudo data.

In this balanced design with extreme small sample sizes, an extreme variance heterogeneity can be seen for higher doses in Figure 4. A modified Dunnett-type test for ratios-to-control allowing heterogeneous variance is used therefore (7).

The fold changes are well reproduced from Tab. 5 of (20), but the Dunnett test for ratio-to-control (assuming normal distributed errors with any variance) reveals a weak significant increase for the 15 mg dose only (adjusted *p*-value) – primarily induced by this extreme small variance, i.e. the three similar values such as (18.7, 17.3, 18.6) by a value range from about 7.2 to 72.4. There is no biological sound increase at any dose – induced by *n*_*i*_ = 3 and the extreme variances at high doses. The conclusion by the authors *”Urinary KIM-1 excretion was increased at doses between 10 and 30mgkg, but owing to a high group variance (i.e. large SDs), the increases of 3.31- and 4.10-fold at, respectively, 45 and 90mgkg HCBD, compared with the control group, were not statistically significant”* can not be reproduced. This example shows very nicely what the *p*-values would have looked like if slightly higher sample sizes had been used, e.g. *n*_*i*_ = 6, just running the simulation code with *n* = 6, assuming that the higher sample size would have the same means and SD.

The well-known fact that the sample size has a substantial influence on the *p*-value becomes very clear, which is particularly important in toxicology.

### 3.3 Taking variance heterogeneity into account

The common assumption of homogeneous variances in dose-responses relationships may be sometimes unrealistic. Corresponding tests tend to be liberal when higher variances occur in higher doses, particularly when sample sizes are smaller in the dose groups (vice versa). In these cases, a modified Dunnett-type test allowing heterogeneous variance should be used (7). As an example we use summary data of pregnant mice which were exposed to five phthalates and their mixture. In their Table 2 (3), the body weight mean, SD and *n* for each treatment group is reported together with stars for Dunnett’s post hoc test. From the simulated data in Figure 5 an unbalanced design with heterogeneous variance can be seen.

**Figure 5:**
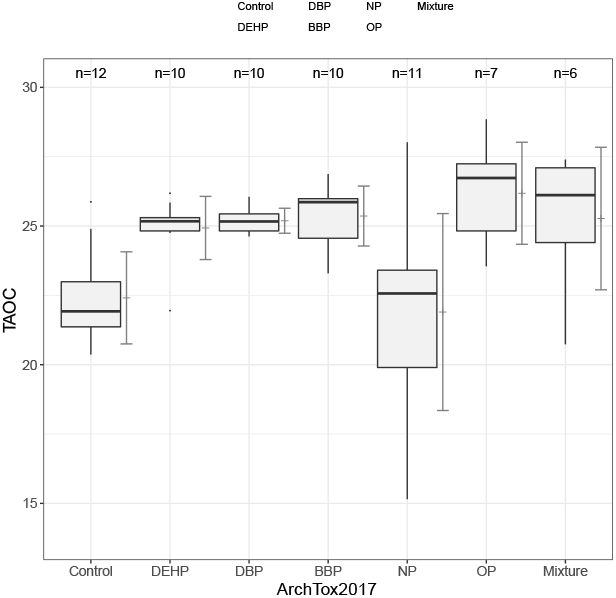
Box plot body weights.

Consequently, the re-analysis with the Dunnett test assuming homogeneous variance reproduced the significance signs. But taking variance heterogeneity into account, there is no significant body weight change in the mixture, but more significant changes in the phthalate groups. This is not supported by the conclusion by the authors *Data showed that, except for NP, all mice exposed to EDCs, single or mixed, presented higher BW than control mice*.

### 3.4 Considering dose as quantitative covariate and/or qualitative factor

Trend tests are classified into those that accept the dose as a factor, i.e. qualitative, and those that accept the dose as a covariate, i.e. quantitative. Both concepts have advantages and disadvantages, but they can even be combined (15). As an example the clinical chemistry endpoints of a feeding study with GMO-corn for the female rats, where the endpoint CHO was selected (Table 4) (5).

**Table 4:** Ratio-to-control inference for *n* = 3.

|  | Fold change | p-value |
| --- | --- | --- |
| 5/0 | 0.97 | 1.00 |
| 10/0 | 1.37 | 0.20 |
| 15/0 | 1.96 | 0.04 |
| 20/0 | 2.39 | 0.17 |
| 30/0 | 4.92 | 0.31 |
| 45/0 | 3.31 | 0.71 |
| 90/0 | 4.10 | 0.59 |

**Table 5:** Ratio-to-control inference for *n* = 6.

| | Fold change | \$p\$-value |
| --- | --- | --- |
| 5/0 | 0.97 | 1.000 |
| 10/0 | 1.37 | 0.015 |
| 15/0 | 1.96 | 0.000 |
| 20/0 | 2.39 | 0.011 |
| 30/0 | 4.92 | 0.051 |
| 45/0 | 3.31 | 0.405 |
| 90/0 | 4.10 | 0.260 |

Assuming the concentration as a quantitative covariate, no significant increasing trend reveals. However, taking additionally the Williams contrast with dose as factor into account, a clear significant plateau effect (contrast C3) follows. This finding is more precise than the significance signs for the 12.5 and 50 group in Table 4 of (5).

### 3.5 User-defined comparisons

Sometimes in trend analysis, additional comparisons than against control are of interest, e.g. control versus high dose (11). As an example, the endpoint VAP (average path velocity) in the gestational exposure to Bisphenol-A affects the spermatozoa in adult mice (12).

**Table 6:**
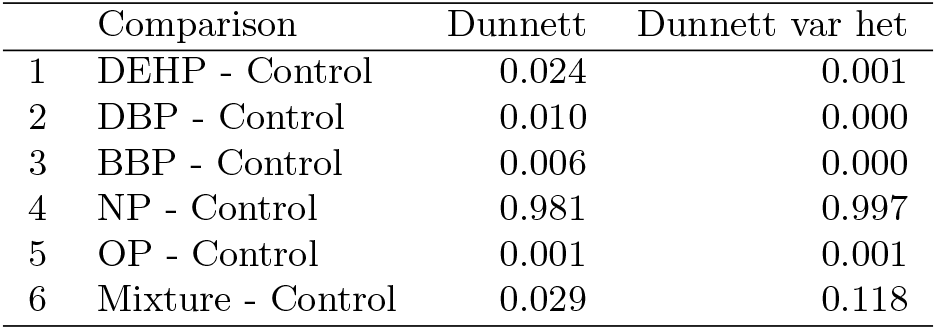
Modified Dunnett test allowing heterogeneous variances

**Table 7:** Tukey and Williams Trend Test

| Dose metameter | Test statistic | *p*-value |
| --- | --- | --- |
| Doseari: Doseari | 1.2759 | 0.2471 |
| Doseord: Doseord | 1.4505 | 0.1898 |
| Doselog: Doselog | -2.4577 | 0.9999 |
| Dosearilog: Dosearilog | 1.4505 | 0.1897 |
| Dosetreat: C 1 | 3.5488 | 0.0007 |
| Dosetreat: C 2 | 3.2176 | 0.0023 |
| Dosetreat: C 3 | 4.7718 | 0.0000 |

**Table 8:** User-defined comparisons

|  | Comparison | Test statistic | *p*-value |
| --- | --- | --- | --- |
| Trend with arithmetic scores | Doseari: Doseari | -3.346 | 0.001 |
| Trend with ordinal scores | Doseord: Doseord | -3.067 | 0.002 |
| Trend with logarithmic scores | Dosearilog: Dosearilog | -2.073 | 0.042 |
| High dose vs. control | Dosetreat: 1 | -4.778 | 0.000 |

Just to report a significant trend test is one story, a different one is to demonstrate a significant trend and a high dose vs. control test (11).

### 3.6 Multiple endpoints

Although multiple endpoints occur frequently in long-term bioassays, such as hematology, commonly they are analysed univariately and independently. A simultaneous test over correlated endpoints can be formulated, to avoid a too high false positive decision rate (8). As an example, the changes in the activities of GSH-R levels in the testicular tissues of male rabbits after treatment with a daily dose of four nitrosamines (MEN, DEN, DMN, DPN) (17) are used. In their Table 1, the bivariate data set of GSH content and GSH-R activity was selected, assuming a correlation of 0.9 (no such correlation information available).

**Table 9:** Bivariate Dunnett Test

| Comparisons | Test statistic | *p*-value |
| --- | --- | --- |
| HL: den - con | -23.3114 | 0.0000 |
| HL: dmn - con | -18.3434 | 0.0000 |
| HL: dpn - con | -21.9466 | 0.0000 |
| HL: men - con | -31.0091 | 0.0000 |
| HR: den - con | -28.5824 | 0.0000 |
| HR: dmn - con | -17.8634 | 0.0000 |
| HR: dpn - con | -27.0118 | 0.0000 |
| HR: men - con | -16.2251 | 0.0000 |

Seriously decreasing levels for both GSH content and GSH-R activity in each of the nitrosamine compounds against control exists under the assumption of a correlated bivariate normal distribution.

### 3.7 A counterexample where normal distribution assumption is unrealistic

Many endpoints in toxicology are not continuous, they are discrete such as counts, e.g., number of micronuclei or proportions, such as tumor rates. For these discrete endpoints the limitation of the above approach based on the normal distribution assumption is obvious. As an example the increase of mutant frequency (MF) after in utero exposure to three doses of benzopyrene in bone marrow (12) (their Figure 3) is considered where the appendix contains the summary statistics. Clearly, MF is a count and was consequently analysed in the generalized linear model in that paper assuming overdispersed Poisson counts. Although counts can be generated as well by the above approach, the limitation of normal distribution assumption becomes obvious in Figure 6 A with jittered box plots, whereas for the 20 and 40 mg group the raw data can be estimated from this figure. This makes a difference in re-analysis, see the boxplots and the evaluation for either Tukey-type trend test or Dunnett-type comparisons against control. The *p*-values for simulated data from mean and SD are unrealistic tiny for any comparisons, whereas ones based on estimated data are more realistic. This highlights the limitation of the ubiquitous use of arithmetic mean and SD as summary statistics – irrespective of the underlying distribution and shows the advantage of raw data presentation within boxplots (13). Comparing the the odds ratios to control (and their simultaneous confidence intervals) for both data sets, reveals a false positive tendency in the simulated data set (see appendix).

**Figure 6:**
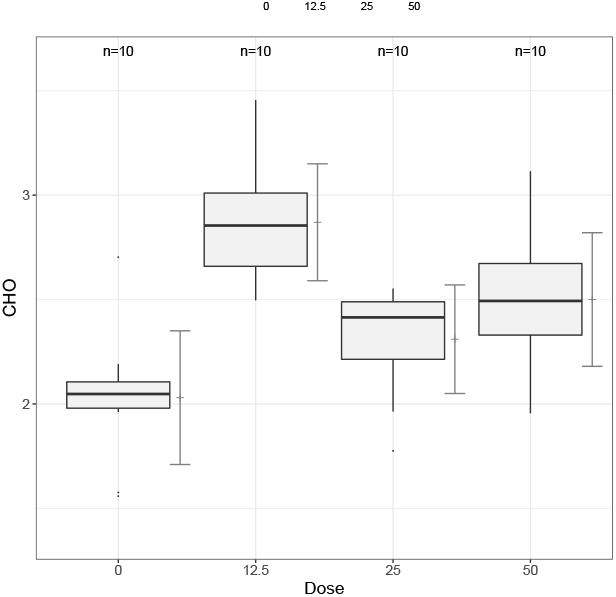
Boxplot CHO endpoint in female rats.

**Figure 7:**
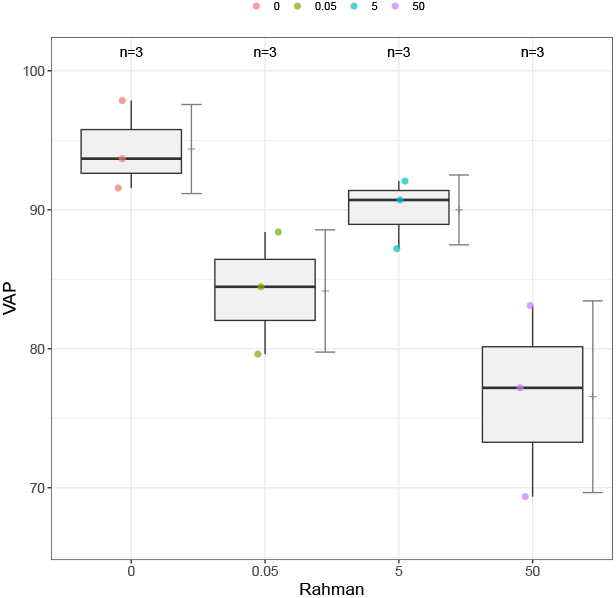
Boxplot VAP.

**Figure 8:**
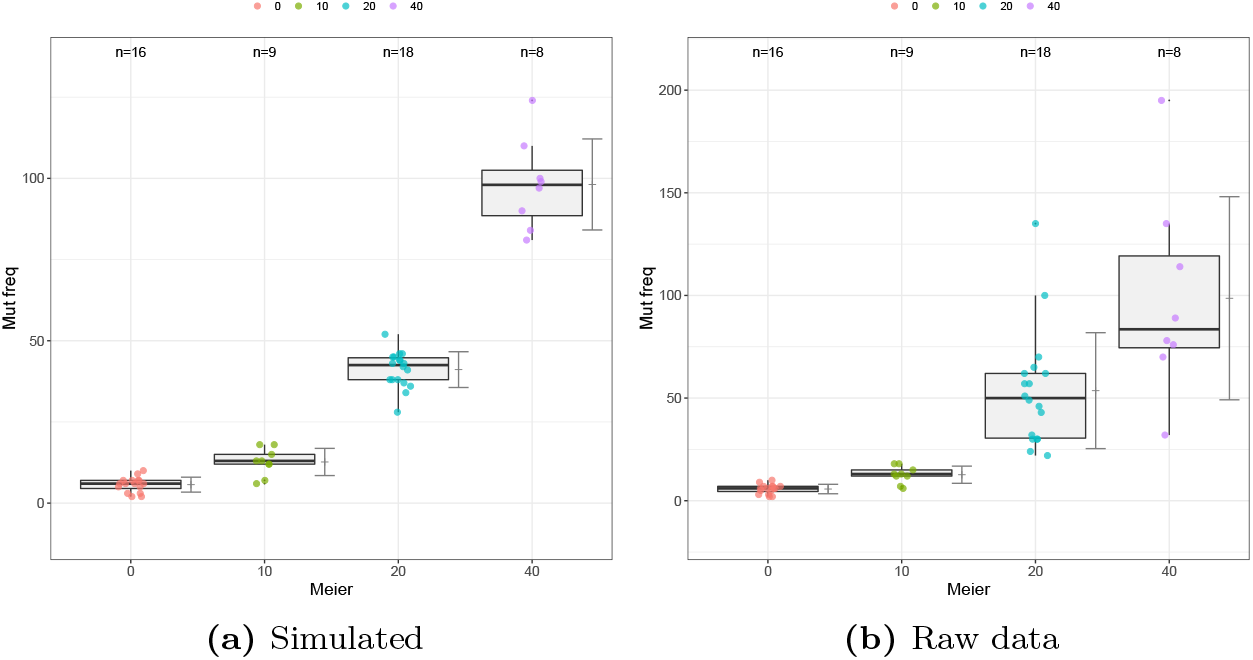
Box bone marrow.

## 4 Discussion

Based on published summary data (means, standard deviations, sample sizes) simulation-based pseudo-data can be used for statistical re-analysis. Examples are: i) Dunnett-type testing instead of independent pairwise t-tests, e.g. for NOAEL identification; ii) test statistics allowing variance heterogeneity, particularly in unbalanced designs; iii) combining multiple bioassay studies, e.g. by using mixed effect models; iv) Analysing trends by alternative models, e.g. using dose as quantitative covariable or a combination of quanti- and qualitative formulation; v) multiple endpoint analysis (instead of independent univariate analysis) assuming low to high correlation between the endpoints; vi) using historical control data in addition to the concurrent control; vii) using robustified trend tests for possible downturn effects at high doses; viii) providing confidence intervals for the effect sites *µ*_*i*_ − *µ*_0_ or *µ*_*i*_/*µ*_0_; ix) using the generated *x*_0,*i*_ as part of a historical control data base; x) simulating their influence of virtual larger or smaller *n*_*i*_ on false negative decision rate; xi) deriving the impact of considering less dose groups, e.g. only 0, 10, 100 instead of 0, 10, 50, 100. Moreover, this approach can be used for teaching purposes, to establish new statistical approaches in toxicology and generating compound/strain/endpoint/sex/dose-specific data base from historical publications on individual data. For the case studies here, the data and R-code is available in the Appendix.

## Acknowledgements

Prof. Dilba, SDSU, for an early version of the algorithm

## 5 Appendix: R-Code

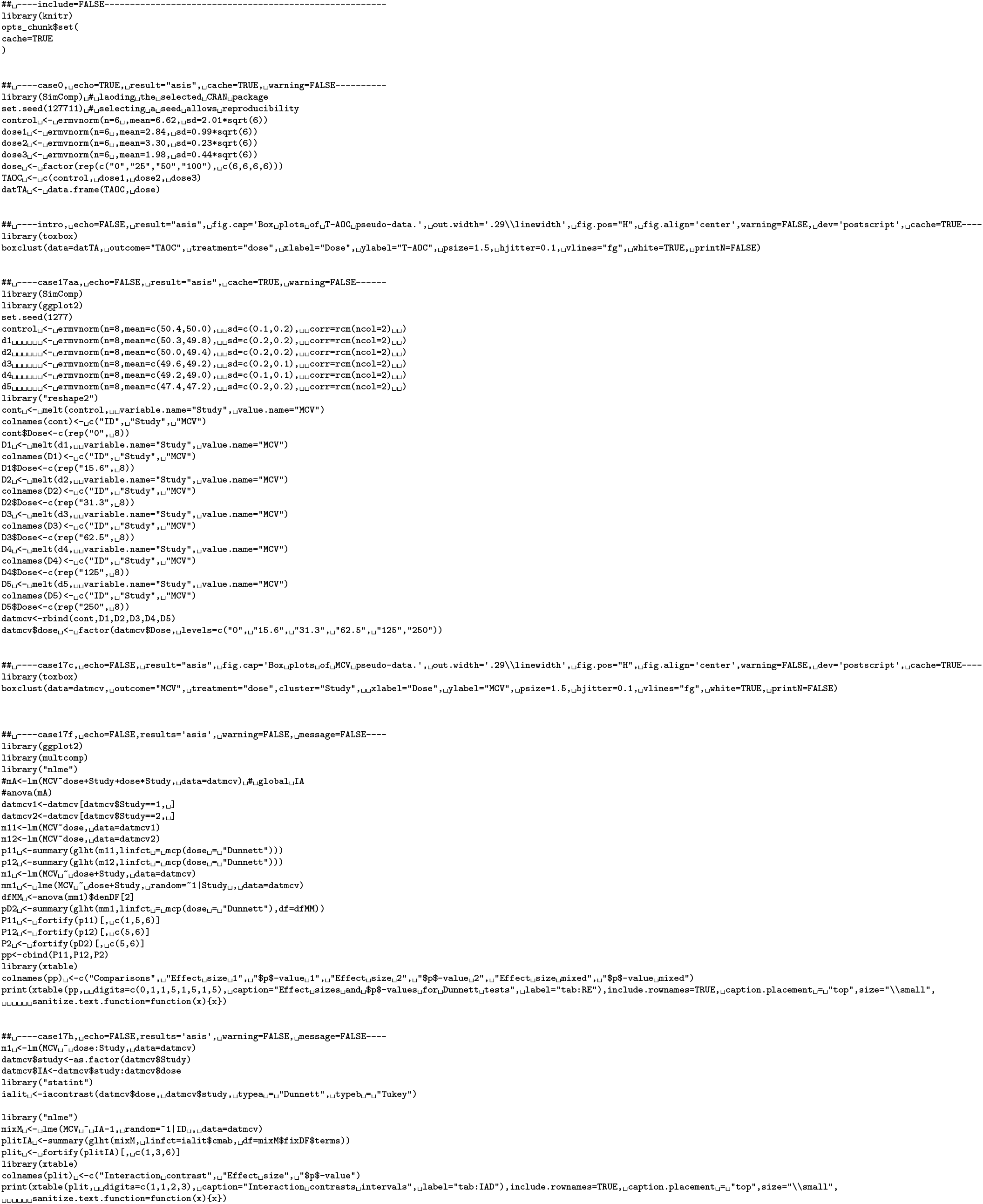
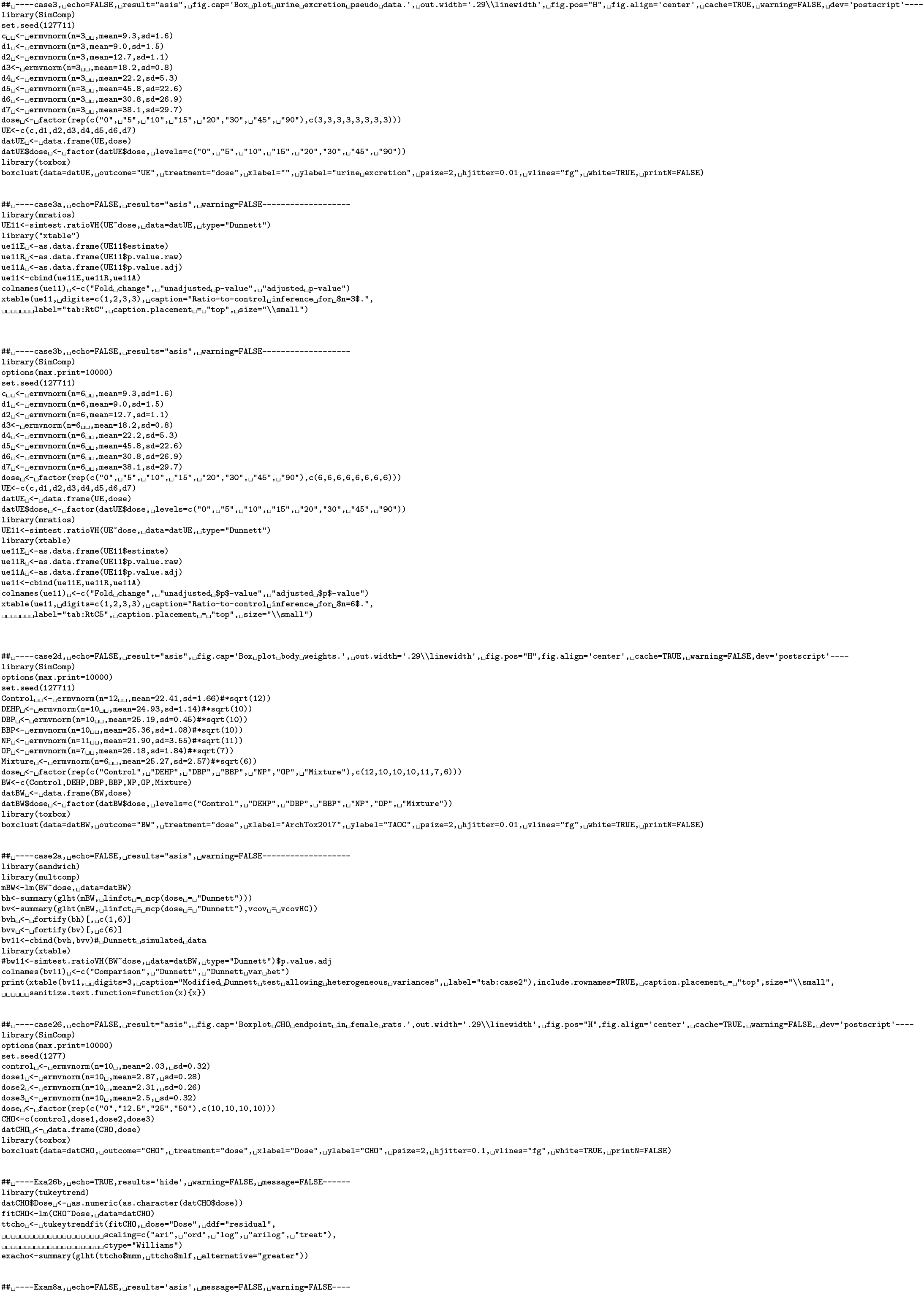
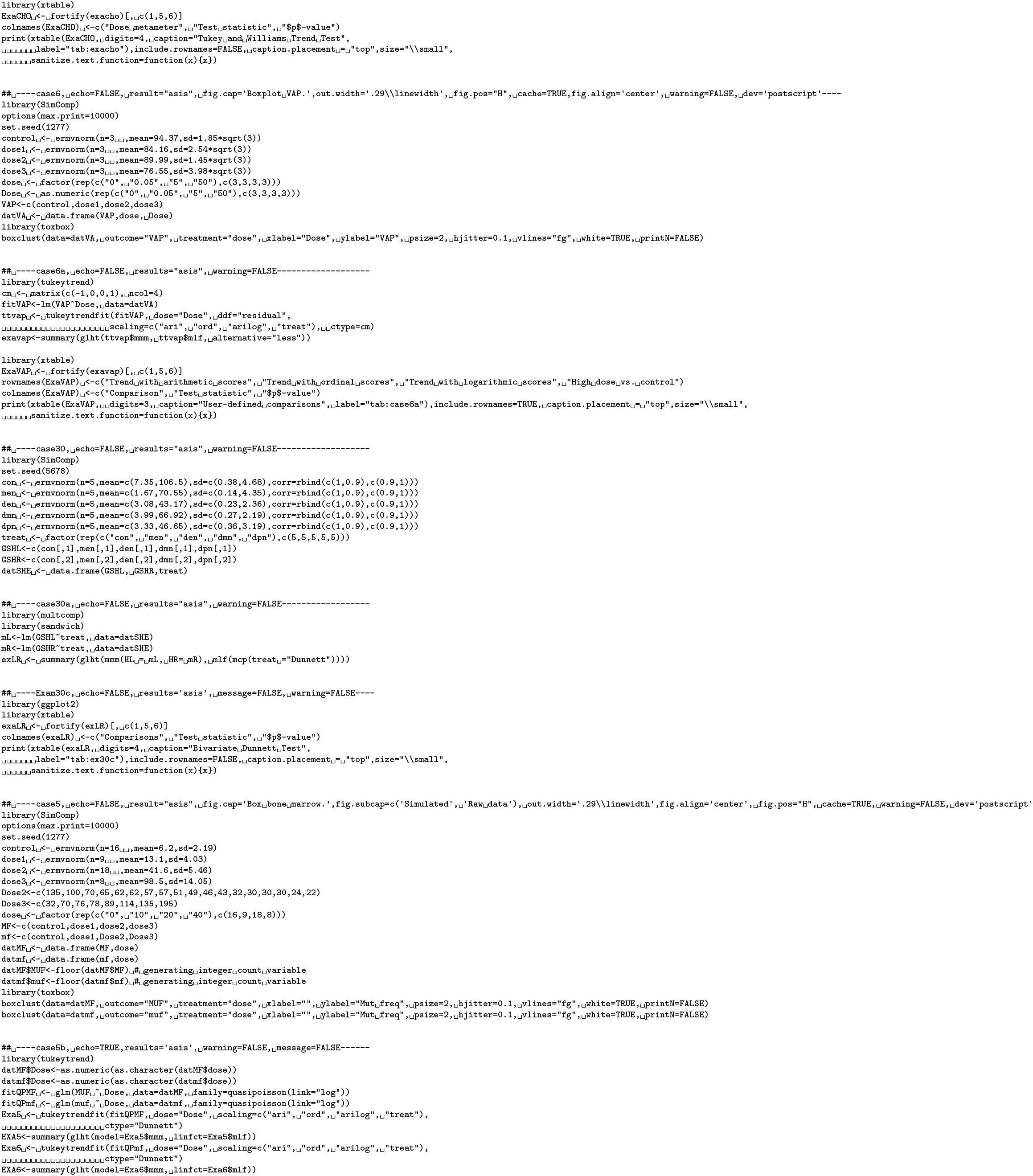

## References

[1] V. Amrhein, S. Greenland, B. McShane, Retire statistical significance, Nature 567 (7748) (2019) 305–307 (Mar. 2019).

[2] [Anonymous]. Kick the bar chart habit. Nature Methods, 11(2):113–113, February 2014.

[3] J. Bunay, E. Larriba, D. Patino-Garcia, L. Cruz-Fernandes, S. Castaneda-Zegarra, M. Rodriguez-Fernandez, J. del Mazo, and R. D. Moreno. Editor’s highlight: Differential effects of exposure to single versus a mixture of endocrine-disrupting chemicals on steroidogenesis pathway in mouse testes. Toxicological Sciences, 161(1):76–86, January 2018.

[4] N. F. Cariello and W. W. Piegorsch. The Ames test: The two-fold rule revisited. Mutation Research-Genetic Toxicology, 369(1-2):23–31, July 1996.

[5] Zhu Y et al. A 90-day feeding study of glyphosate-tolerant maize with the g2-aroa gene in sprague-dawley rats. Food Chem Tox, 2013.

[6] A. J. Fosang and R. J. Colbran. Transparency is the key to quality. Journal of Biological Chemistry, 290(50):29692–29694, December 2015.

[7] M. Hasler and L. A. Hothorn. Multiple contrast tests in the presence of heteroscedasticity. Biometrical Journal, 50(5):793–800, October 2008.

[8] M. Hasler and L. A. Hothorn. Multi-arm trials with multiple primary endpoints and missing values. Statistics in Medicine, 37(5):710–721, February 2018.

[9] L. A. Hothorn and G. D. Djira. A ratio-to-control williams-type test for trend. Pharmaceutical Statistics, 10(4):289–292, July 2011.

[10] A. Kitsche and L. A. Hothorn. Testing for qualitative interaction using ratios of treatment differences. Statistics in Medicine, 33(9):1477–1489, April 2014.

[11] K. K. Lin and M. A. Rahman. Comparisons of false negative rates from a trend test alone and from a trend test jointly with a control-high groups pairwise test in the determination of the carcinogenicity of new drugs. J. Biopharm Statist, 2018.

[12] M. J. Meier, J. M. O’Brien, M. A. Beal, B. Allan, C. L. Yauk, and F. Marchetti. In utero exposure to benzo[a]pyrene increases mutation burden in the soma and sperm of adult mice. Environmental Health Perspectives, 125(1):82–88, January 2017.

[13] P. Pallmann and L. A. Hothorn. Boxplots for grouped and clustered data in toxicology. Archives of Toxicology, 90(7):1631–1638, July 2016.

[14] B.D. Ripley. Stochastic simulation. Wiley, 1987.

[15] Schaarschmidt. Multiplicity adjustment for tukeys trend test and extensions to the multiple marginal generalized linear and linear mixed models.

[16] F. Schaarschmidt. Simultaneous confidence intervals for multiple comparisons among expected values of log-normal variables. Computational Statistics and Data Analysis, 58:265–275, FEB 2013.

[17] S. A. Sheweita, Y. Y. El Banna, M. Balbaa, I. A. Abdullah, and H. E. Hassan. N-nitrosamines induced infertility and hepatotoxicity in male rabbits. Environmental Toxicology, 32(9):2212–2220, September 2017.

[18] K. A. Shipkowski, C. M. Sheth, M. J. Smith, M. J. Hooth, K. L. White, and D. R. Germolec. Assessment of immunotoxicity in female fischer 344/n and sprague dawley rats and female b6c3f1 mice exposed to hexavalent chromium via the drinking water. Journal of Immunotoxicology, 14(1):215–227, November 2017.

[19] Z. L. Sun and S. J. et al.. Li. Alterations in epididymal proteomics and antioxidant activity of mice exposed to fluoride. Archives of Toxicology, 92(1):169–180, January 2018.

[20] A. Swain, J. Turton, C. Scudamore, D. Maguire, I. Pereira, S. Freitas, R. Smyth, M. Munday, C. Stamp, M. Gandhi, S. Sondh, H. Ashall, I. Francis, J. Woodfine, J. Bowles, and M. York. Nephrotoxicity of hexachloro-1:3-butadiene in the male hanover wistar rat; correlation of minimal histopathological changes with biomarkers of renal injury. Journal of Applied Toxicology, 32(6):417–428, June 2012.

[21] T. L. Weissgerber, V. D. Garovic, S. J. Winham, N. M. Milic, and E. M. Prager. Transparent reporting for reproducible science. Journal of Neuroscience Research, 94(10):859–864, October 2016.

